# Maintenance of Cell Fates by Regulation of the Histone Variant H3.3 in *Caenorhabditis elegans*

**DOI:** 10.1101/294827

**Authors:** Yukimasa Shibata, Kiyoji Nishiwaki

## Abstract

**Highlights:** TLK-1 maintains cell fates by repression of selector genes

TLK-1 and downstream H3 chaperone CAF1 inhibit H3.3 deposition

Loss of *sin-3* suppresses the defect in cell-fate maintenance of *tlk-1* mutants

AcH4-binding protein BET-1 is necessary for *sin-3* suppression

**Summary:** Cell-fate maintenance is important to preserve the variety of cell types that are essential for the formation and function of tissues. We previously showed that the acetylated histone H4-binding protein BET-1 maintains cell fate by recruiting the histone variant H2A.z. Here, we report that *Caenorhabditis elegans* tousled-like kinase TLK-1 and the histone H3 chaperone CAF1 maintain cell fate by preventing the incorporation of histone variant H3.3 into nucleosomes, thereby repressing ectopic expression of transcription factors that induce cell-fate specification. Genetic analyses suggested that TLK-1 and BET-1 act in parallel pathways. In *tlk-1* mutants, the loss of SIN-3, which promotes histone acetylation, suppressed a defect in cell-fate maintenance in a manner dependent on MYST family histone acetyltransferase MYS-2 and BET-1. *sin-3* mutation also suppressed abnormal H3.3 incorporation. Thus, we propose that the regulation and interaction of histone variants play crucial roles in cell-fate maintenance through the regulation of selector genes.

## Introduction

Defects in cell-fate maintenance cause aberrant cell-fate transformation, which can induce tumor formation and tissue malfunction. Conversely, suppression of the mechanisms that maintain cell fate is necessary for efficient reprogramming such as the generation of induced pluripotent stem (iPS) cells (Takahashi and Yamanaka, 2015). Aberrant activation of genes that induce specific cell fates causes abnormal cell-fate transformation (Halder et al., 1995; Riddle et al., 2013). Thus, the repression of the genes that specify cell fates is critical for maintaining individual cell fates.

Epigenetic marks including histone modifications play important roles in transcriptional repression during development. For example, methylation on lysine 27 of histone H3 (H3K27me) is required to silence developmentally regulated genes such as Hox genes (Ringrose and Paro, 2004). In contrast, the roles of histone variants in transcriptional repression are poorly understood. We previously showed that a histone H2A variant, H2A.z, is required to maintain cell fate in multiple cell lineages in *Caenorhabditis elegans* (Shibata et al., 2014). Subnuclear localization of H2A.z is regulated by an acetylated histone H4– binding protein, BET-1, that is also required to maintain cell fate. BET-1 represses selector genes that encode DNA-binding transcription factors (TFs) such as LIM homeodomain protein MEC-3 and CEH-22/Nkx2.5, which induce specific cell fates (Shibata and Nishiwaki, 2014; Shibata et al., 2014; Shibata et al., 2010). The selector gene activates transcription of itself and of genes that are required for the specific function of each cell type (Hobert, 2008). Thus, although many studies suggest a role for H2A.z in transcriptional activation, H2A.z also preserves transcriptional repression in the maintenance of cell fate.

In addition to the H2A variant, another major histone variant is the histone H3 variant H3.3, which is often observed on actively transcribed loci (Wirbelauer et al., 2005). Canonical histone H3 and H3 variant H3.3 are deposited by chromatin assembly factor 1 (CAF1) and histone regulator A (HIRA), respectively (Tagami et al., 2004). In cultured cells, CAF1 depletion causes alternative deposition of H3.3 to fill the nucleosome gap at the replication site by HIRA (Ray-Gallet et al., 2011). CAF1 deficiency promotes artificial trans-differentiation, such as induction of iPS cells and the generation of neurons from fibroblasts and of macrophages from pre-B cells (Cheloufi et al., 2015). However, the roles of CAF1 and H3.3 in cell-fate maintenance during development are not known.

Several studies have suggested a relationship between H2A.z and H3.3. Genome-wide analyses of H2A.z, RNA pol II, and transcription suggest that H2A.z is correlated with transcriptional repression at the poised state (Raisner et al., 2005). Biophysical evidence indicates that H2A.z promotes chromatin compaction (Chen et al., 2013). In contrast, H3.3 promotes gene activation, counteracting H2A.z-mediated transcriptional repression, and impairs H2A.z-mediated chromatin compaction, thus reducing higher-order chromatin folding (Chen et al., 2013). Because transcriptional repression by H2A.z is necessary to maintain cell fate, it is important to clarify the role of H3 variants and their regulation to understand the maintenance of cell identity.

Tousled-like kinases (TLKs) are conserved protein kinases in multicellular organisms. They phosphorylate anti-silencing factor 1 (ASF1), which interacts with CAF1 (Klimovskaia et al., 2014). Arabidopsis TLK, Tousled, acts in the maintenance of transcriptional gene silencing and is required for leaf and flower development (Roe et al., 1993; Wang et al., 2007). In *C. elegans* early embryos, the ortholog TLK-1 is required for chromosome segregation and cytokinesis and promotes transcription (Han et al., 2005; Han et al., 2003; Yeh et al., 2010). Our genetic screening for mutants that are defective in cell-fate maintenance resulted in the isolation of *tlk-1* mutants. Here, we analyzed the roles of TLK-1 and CAF1 in cell-fate maintenance and the regulation of H3.3. We also investigated the relationship between mechanisms that regulate H3.3 and H2A.z.

## Results

### Isolation of *tlk-1* mutants by screening for aberrant cell-fate transformation mutants

We previously showed that, in *C. elegans*, malfunction of the machinery that maintains cell fate induces the production of extra distal tip cells (DTCs) (Shibata et al., 2010). In wild-type animals, there are two DTCs that function as leader cells during gonad formation. To identify additional genes that are required for the maintenance of cell fate, we screened for mutants that have extra DTCs and isolated two mutants of *tlk-1* (Fig. 1A–E, Fig. S1A). *tlk-1* encodes a serine/threonine kinase that is a member of the TLK family (Fig. 1F–H). Although humans and mice have two TLK family proteins, TLK-1 is the sole family member in *C. elegans*. *tlk-1*(*tk158)* has a nonsense mutation at Q44stop, and *tlk-1*(*tk170)* has a missense mutation at T846I (Fig. 1H, Fig. S1B). The translational termination near the N terminus suggests that *tk158* is a null allele. A DNA fragment containing the coding region and 3.5 kb of upstream sequence fully rescued the *tlk-1(tk158)* mutant phenotype (Fig. 1E, Fig. S1A). *tlk-1::gfp* expression was observed in the nuclei of all somatic cells including cells of the somatic gonad, neurons in the posterior lateral ganglia (PLG), and the hypodermis (Fig. 1I-L). TLK-1 is also expressed in the nuclei of embryos (Han et al., 2003). The knockdown of *tlk-1* by feeding RNAi resulted in embryonic lethality (data not shown). However, we observed the postembryonic extra-DTC phenotype in *tlk-1* homozygous mutants from heterozygous mutant hermaphrodites because the embryonic lethality was rescued by the maternal effect.

**Fig. 1.**
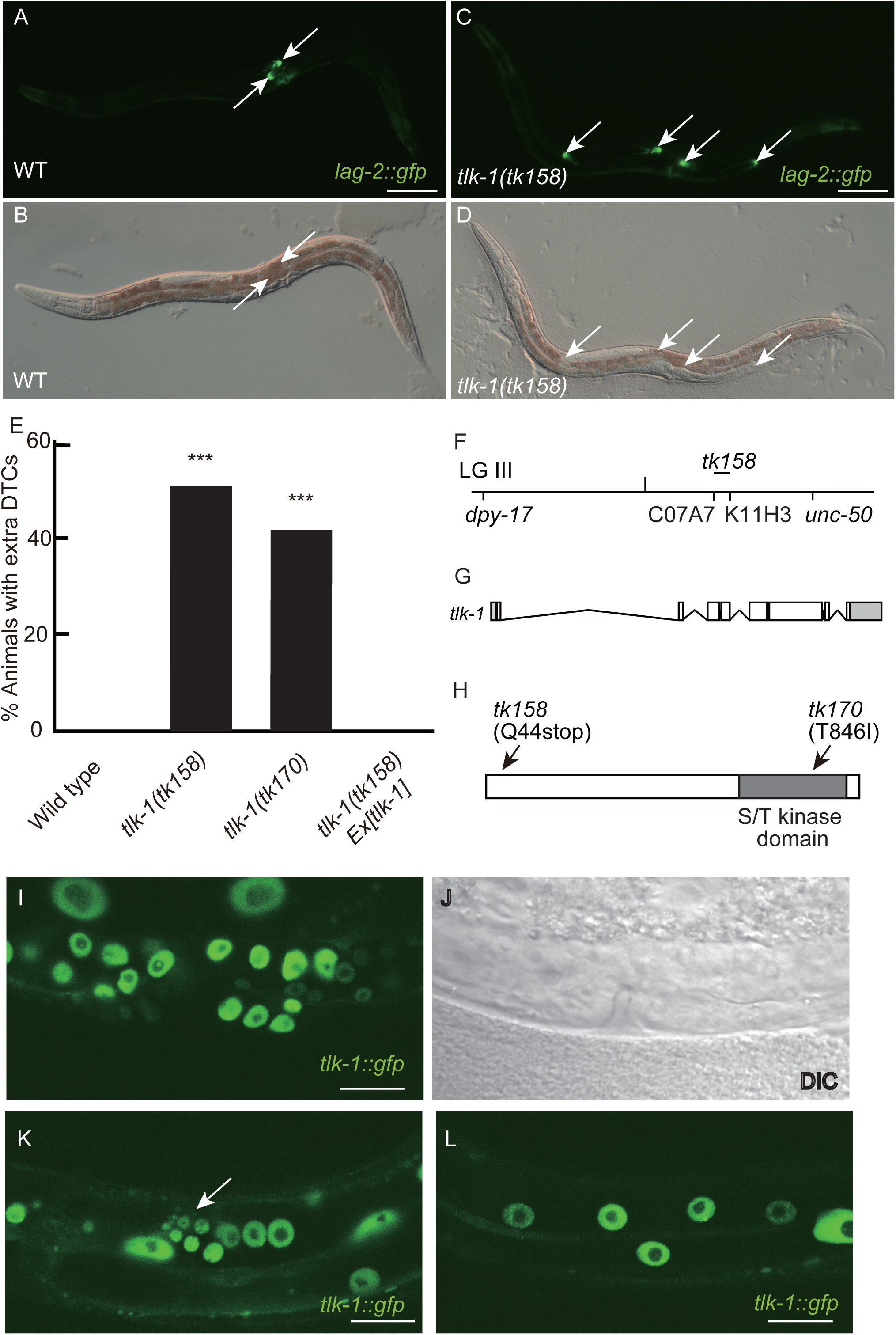
Isolation of *tlk-1* mutants that show the extra-DTC phenotype. (A–D) GFP (A, C) and differential interference contrast (DIC) (B, D) images showing the expression of the DTC marker *lag-2::gfp* in wild type (WT) (A, B) and *tlk-1* mutants (C, D) at the adult stage. Anterior is to the left, ventral is to the bottom. Arrows indicate *lag-2::gfp*-positive cells. Scale bars in A and C indicate 100 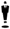 m in A-D. (E) Bar graph showing the percent of adult animals with the extra-DTC phenotype. *n* = 100; \*\*\**p* < 0.005, as compared with wild type. (F–H) Schematic diagrams of the chromosomal position of *tlk-1* (F), *tlk-1* gene structure (G), and TLK-1 protein structure (H). (G) Coding and non-coding regions are indicated by white and gray boxes, respectively. (H) Serine threonine (S/T) kinase domain is indicated by the gray box. Arrows indicate the positions of *tk158* and *tk170* mutations. (I, K, L) *tlk-1::gfp* expression in the somatic gonad (I), PLG (K), and hypodermis (L) of L4 larvae. Arrow indicates the PLG. Scale bars in I, K, and L indicate 10 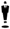 m in I-L. (J) DIC image that corresponds to panel (I).

### TLK-1 regulates cell fate in multiple cell lineages

In the wild-type somatic gonad, the two DTCs express *lag-2::gfp* (Kostic et al., 2003). Extra DTCs were observed in half of the *tk158* mutants (Fig. 1E). The maximum number of DTCs was five cells in *tk158* mutants (Fig. S1A). In addition to expressing *lag-2::gfp*, DTCs were positioned at the tip of the gonad arms and showed a cup-like shape in wild-type animals (Fig. 2A). *tlk-1* mutants had extra DTCs at the tips of the extra gonad arms. The extra DTCs were also cup shaped (Fig. 2B), suggesting differentiation into DTCs rather than simple ectopic expression of *lag-2::gfp*. We examined whether extra DTC formation depended on the NK-2 family homeodomain DNA-binding TF CEH-22, which induces DTCs (Shibata et al., 2014). Because *ceh-22* is required for the production of mother cells of DTCs (Lam et al., 2006), we performed partial knockdown of *ceh-22* by feeding RNAi. *ceh-22* RNAi in *tlk-1* mutants partially suppressed the extra-DTC phenotype (Fig. 2C, Fig. S2A). Therefore, inappropriate expression of *ceh-22* induced extra DTCs in *tlk-1* mutants.

**Fig. 2.**
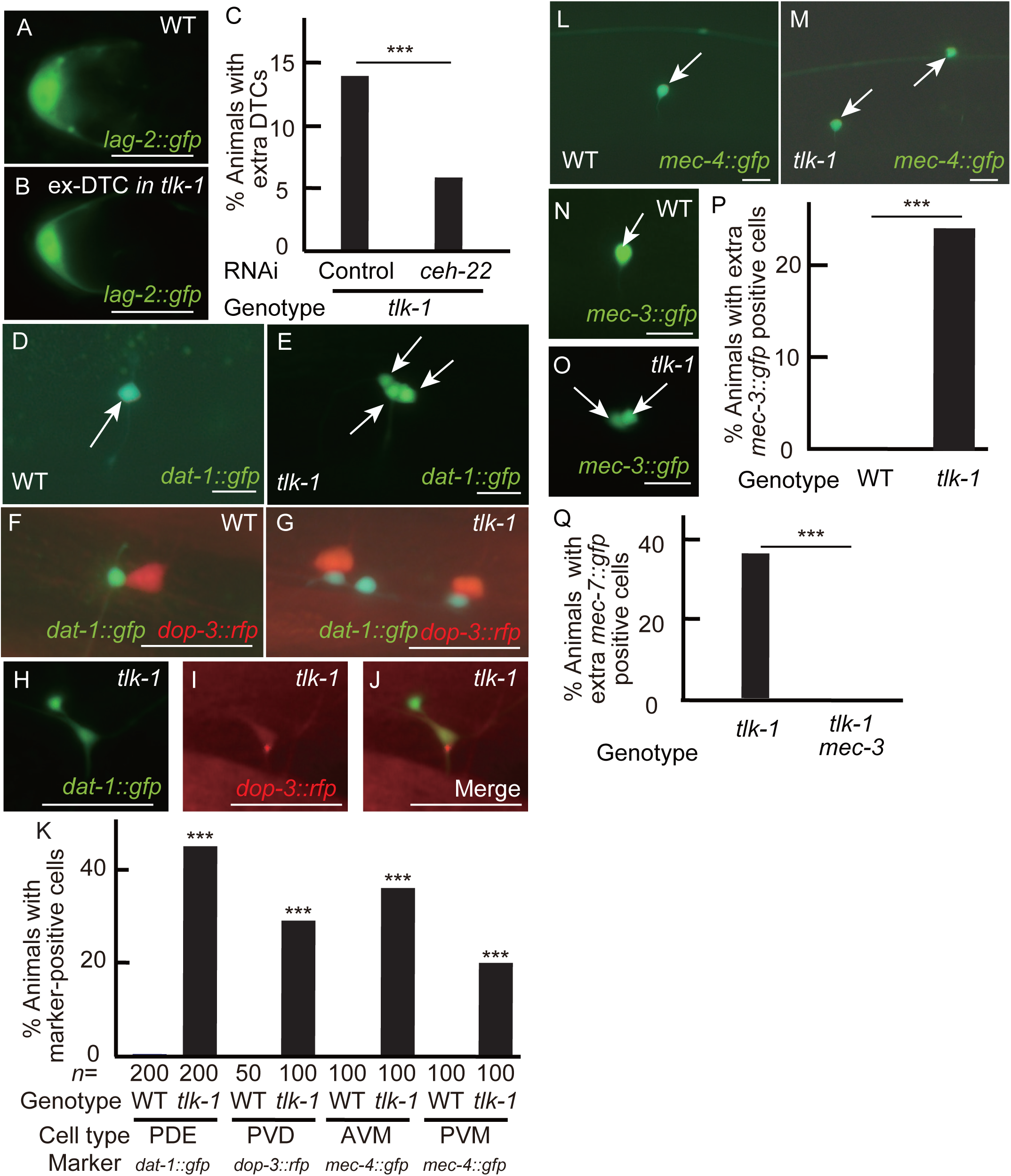
Cell-fate transformation in *tlk-1* mutants. (A, B, D–J, L–O) Fluorescence images showing the expression of markers *lag-2::gfp* (A, B), *dat-1::gfp* (D, E, H), *dat-1::gfp* and *dop-3::rfp* (F, G, J), *dop-3::rfp* (I), *mec-4::gfp* (L, M), and *mec-3::gfp* (N, O) in wild type (WT) (A, D, F, L, N) and *tlk-1* mutants (B, E, G–J, M, O) at the adult stage. Arrows indicate marker-positive cells. (D–J, L–O) Anterior is to the left, ventral is to the bottom. Scale bars indicate 10 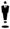 m. (C, K, P, Q) Bar graphs show the percent of adult animals with extra DTCs (C), extra marker-positive cells (K), extra *mec-3::gfp*-positive cells (P), the extra *mec-4::gfp*-positive cells (Q). *n* = 150, 135, and 100 in control of panel C, in *ceh-22* RNAi of panel C, and in panels P and Q, respectively; \*\*\**p* < 0.005.

We next examined whether *tlk-1* functions in other cell types. *dat-1::gfp* and *dop-3::rfp* are expressed in bilateral pairs of PDE and PVD neurons, respectively, in the wild-type PLG (Chase et al., 2004; Nass et al., 2002). *dat-1::gfp* and *dop-3::rfp* markers were ectopically expressed in *tlk-1* mutants in the PLG region (Fig. 2D–G, K, Fig. S2B). We rarely observed cells with both markers; only 3% of *tlk-1* mutants exhibited this phenotype (Fig. 2H–J). In addition to gene expression, we used cell size as a characteristic of cell type. In wild type, PVD (*dop-3::rfp* positive) is larger than other neuronal cells including PDE (*dat-1::gfp* positive) (Fig. 2F). *tlk-1* mutants have multiple *dat-1::gfp*-positive cells and/or *dop-3::rfp*-positive cells, although the number of these cells varies among individuals. As observed in wild-type animals, *dop-3::rfp*-positive cells were larger than *dat-1::gfp*-positive cells in *tlk-1* mutants (Fig. 2G), suggesting differentiation into PDE or PVD cells rather than simple ectopic expression of *dat-1::gfp* or *dop-3::rfp*. Ectopic expression of another PDE marker, *osm-6::gfp*, was also observed in *tlk-1* mutants (Fig. 3H). Thus, ectopic PDE and PVD cells were produced in *tlk-1* mutants.

**Fig. 3.**
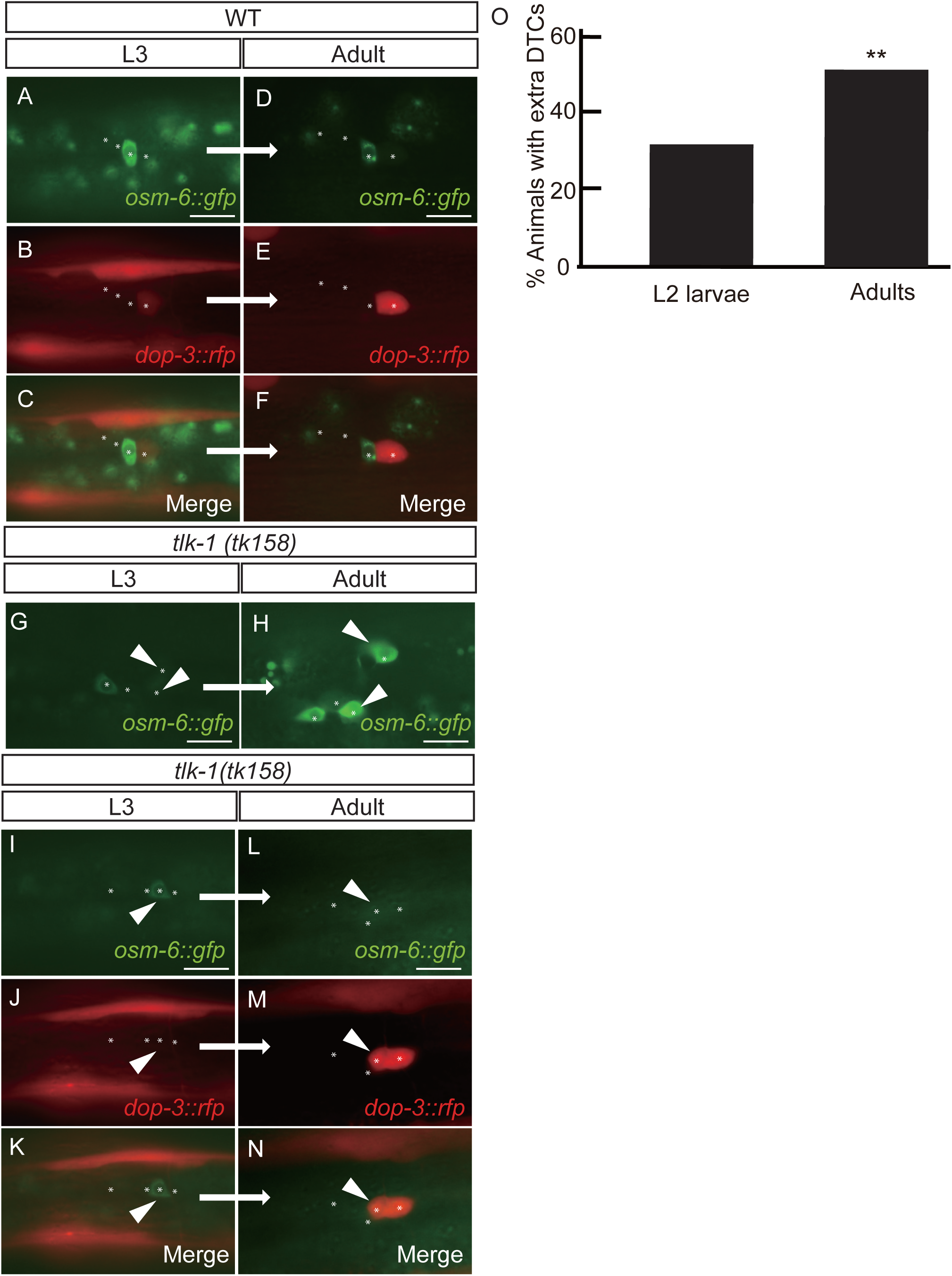
Transformation of cell identity in *tlk-1* mutants. Fluorescence images of wild type (WT) (A–F) and two individual *tlk-1* mutant animals (G–N). Expression was observed at the L3 stage and then at the adult stage in the same animal. Asterisks indicate the positions of neuronal nuclei detected in the DIC images. Arrowheads indicate the cells with altered marker expression between the two stages. (G, H) Some *osm-6::gfp*-negative cells at the L3 stage later expressed *osm-6::gfp* until the adult stage. (I–N) Some *osm-6::gfp*-expressing cells at the L3 stage lost this expression at the adult stage, and the same cells expressed *dop-3::rfp* at the adult stage. Anterior is to the left, ventral is to the bottom. Scale bars indicate 10 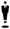 m. (O) Bar graph shows the percent of *tlk-1* mutants with the extra-DTC phenotype among L2 larvae and adults. *n* = 100; **0.005 < *p* < 0.01, as compared with the L2 larvae.

*mec-4::gfp* is expressed in AVM and PVM neurons at the anterior right and posterior left sides, respectively, in wild-type animals (Clark and Chiu, 2003) Ectopic expression of *mec-4::gfp* was observed in the region where wild-type marker-positive cells were observed (Fig. 2L, M, Fig. S2B). In wild-type animals, AVM cells expressed *mec-3* in addition to *mec-4* (Fig. 2N). We found ectopic expression of *mec-3::gfp* in *tlk-1* mutants (Fig. 2O, P, Fig. S2C), suggesting that extra AVMs are produced. *mec-3* encodes a LIM homeodomain DNA-binding TF that induces six mechanosensory neurons including AVM (Way and Chalfie, 1988). *mec-4* is a direct target of MEC-3 (Duggan et al., 1998; Hobert, 2008). If *tlk-1* controls the AVM fate, *mec-4* expression could be regulated by *mec-3*. We examined *mec-4::gfp* expression in *tlk-1 mec-3* double mutants and found no ectopic *mec-4::gfp* expression (Fig. 2Q, Fig. S2D), suggesting that extra AVMs are formed by inappropriate expression of *mec-3* in *tlk-1* mutants. These results suggested that TLK-1 inhibits the production of ectopic cells by repressing genes that encode cell type-specific TFs.

### Cell-fate transformation in *tlk-1* mutants

In wild-type animals, distal granddaughters of the Z1/Z4 cells that are born at the L1 stage differentiate into DTCs until the early L2 stage (Shibata et al., 2010). In *tlk-1* mutants, we compared the extra-DTC phenotype at the L2 and adult stages and found a lower penetrance at the L2 stage (Fig. 3O, Fig. S3A). Analysis of the DTC numbers in the L2 and adult stages in the same animals revealed that they increased in 12 of 29 *tlk-1* mutant animals (data not shown). These results indicated that extra DTCs are produced even after the production of normal DTCs.

To elucidate the cause of extra marker-positive cells, we observed the PLG using the PDE marker *osm-6::gfp* and the PVD marker *dop-3::rfp* at the L3 and adult stages in the same animals. The position of the cells is variable in each animal, but the relative position of cells is conserved in the same animal during development (Shibata et al., 2010). In wild-type animals, cells that expressed *osm-6::gfp* at the L3 stage never expressed *dop-3::rfp* at the adult stage and vice versa (Fig. 3A–F). Cell-fate transformation in the PLG between the L3 stage and the adult stage was not observed in wild-type animals (*n* = 13). In contrast, we found that *osm-6::gfp*-negative cells at the L3 stage (21.5 hr after hatching) started expressing *osm-6::gfp* until the adult stage in 2 of 13 *tlk-1* mutants (Fig. 3G, H). In addition, in 3 of 13 *tlk-1* mutants, *osm-6::gfp*-expressing cells at the L3 stage no longer expressed this marker at the adult stage, and the same cell expressed *dop-3::rfp* (Fig. 3I-N), suggesting the transformation from the PDE to the PVD fate. We also observed the same *tlk-1* mutant animals at the L3 (24 hr after hatching) and adult stages. Of 25 *tlk-1* mutants, 2 expressed both *osm-6::gfp* and *dop-3::rfp* in the same cells at the L3 stage, but these *osm-6::gfp* and *dop-3::rfp* double-positive cells became *dop-3::rfp* single-positive cells at the adult stage (Fig. S3B). Thus, transformation from the PDE to the PVD fate occur at least at the L3 stage. These results indicated that abnormal cell-fate transformation occurs in *tlk-1* mutants and that TLK-1 maintains cell fate in multiple cell lineages.

### Disruption of histone chaperone CAF1 causes a *tlk-1*-like phenotype

TLK phosphorylates ASF1 to promote histone supply to the CAF1 complex, which deposits the histone H3-H4 complex as a DNA replication-coupled histone chaperone (Klimovskaia et al., 2014). *C. elegans* has two ASF1 homologs, UNC-85 and ASFL-1 (Grigsby and Finger, 2008; Grigsby et al., 2009), but neither *unc-85* nor *asfl-1* mutants show the extra-DTC phenotype (Fig. 4E). In addition, *unc-85 asfl-1* double-homozygous mutants from *unc-85* homozygous and *asfl-1* heterozygous (*asfl-1/hT2; unc-85*) hermaphrodites did not show the extra-DTC phenotype. *unc-85 asfl-1* double-homozygous mutants from *unc-85 asfl-1* double-homozygous mutants were embryonic lethal. Thus, there was no evidence to show the requirement of ASF1 homologs in cell-fate maintenance.

**Fig. 4.**
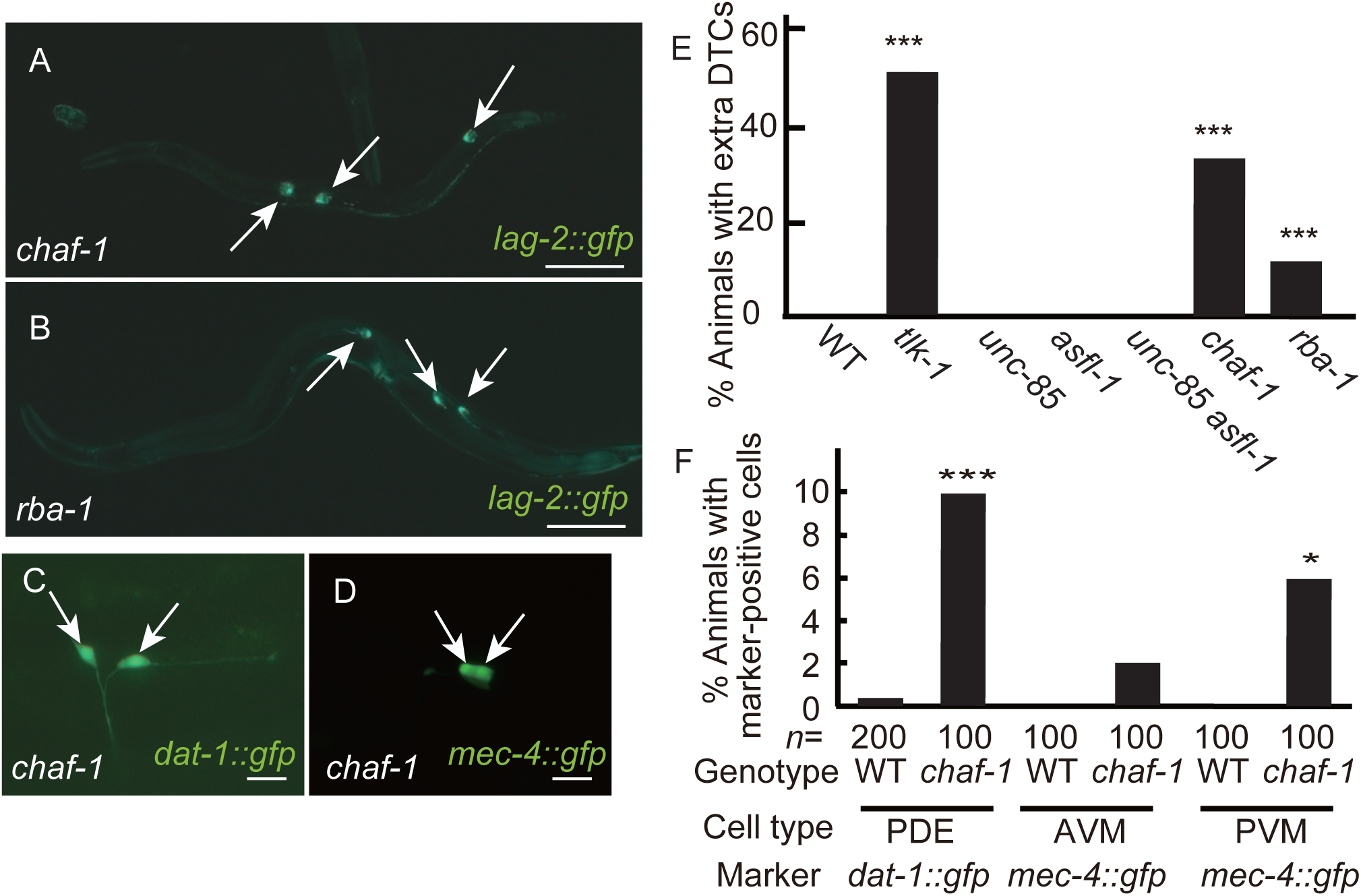
Disruption of the CAF1 complex mimics *tlk-1* mutation. (A–D) GFP images showing the expression of *lag-2::gfp* (A, B), *dat-1::gfp* (C), and *mec-4::gfp* (D) in *chaf-1* (A, C, D) or *rba-1* mutants (B) at the adult stage. Arrows indicate marker-positive cells. Scale bars indicate 100 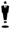 m in (A, B) and 10 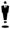 m in (C, D). (E, F) Bar graphs shows the percent of adult animals with the extra-DTC phenotype (E) and with *dat-1::gfp*-positive or *mec-4::gfp*-positive cells (F). (E) *n* = 100; *0.05 < *p* < 0.01, \*\*\**p* < 0.005, as compared with the wild type.

*C. elegans* counterparts of CAF1 subunits are CHAF-1, CHAF-2, and RBA-1 (Nakano et al., 2011). We examined *chaf-1* mutants and *rba-1* mutants and found that they exhibited the extra-DTC phenotype (Fig. 4A, B, E, Fig. S4A). We also examined *dat-1::gfp* and *mec-4::gfp* expression in *chaf-1* mutants. As in *tlk-1* mutants, extra *dat-1::gfp*-positive cells and extra *mec-4::gfp*-positive cells were observed (Fig. 4C, D, F, Fig. S4B).Thus, the CAF1 deficiency caused phenotypes similar to those observed in *tlk-1* mutants. In addition, *tlk-1 chaf-1* double mutants showed more severe phenotypes than did *tlk-1* or *chaf-1* single mutants (Fig. 6A), suggesting that CAF1 activity was down-regulated but not abolished in *tlk-1* mutants.

### Nuclear H3.3 levels are up-regulated in *tlk-1* mutants

In cultured cells, CAF1 depletion causes alternative deposition of H3.3 that is observed in actively transcribed loci (Ray-Gallet et al., 2011; Wirbelauer et al., 2005). To examine the level of H3.3 deposition onto chromatin in *chaf-1* or *tlk-1* mutants, we observed the expression of *his-72*, which encodes H3.3 (Boeck et al., 2011; Ooi et al., 2006). Higher *his-72::gfp* expression was observed in the nuclei of the hypodermis of *tlk-1* and *chaf-1* mutants than in the wild type (Fig. 5A, B, E). The expression level was higher in *tlk-1* mutants than in *chaf-1* mutants, which was consistent with higher penetrance of the extra-DTC phenotype in *tlk-1* mutants. The granular pattern of *his-72::gfp* expression in the nucleus suggested that HIS-72::GFP was deposited in the chromatin. Higher *his-72::gfp* expression was also observed in the somatic gonad of *tlk-1* mutants relative to the wild-type somatic gonad (data not shown). In contrast to *tlk-1* mutants, mutants of *bet-1*, which functions through the deposition of H2A.z (Shibata et al., 2014), showed only weak up-regulation of *his-72::gfp* expression (Fig. 5E). These results indicated that TLK-1 and CAF1 reduce *his-72::gfp* accumulation in chromatin.

**Fig. 5.**
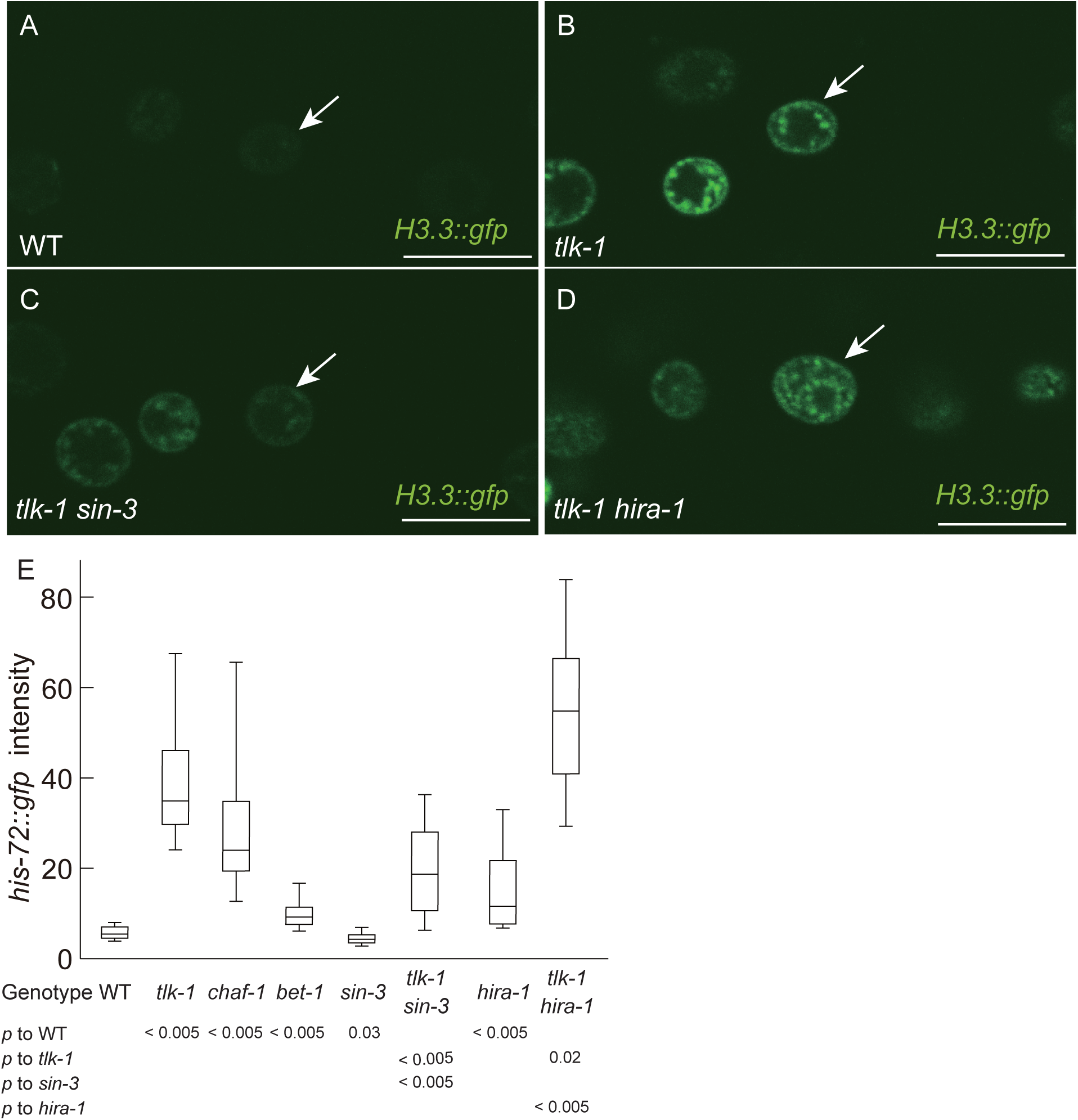
*tlk-1* controls H3.3 amounts in the nucleus. (A–D) GFP images of V5.ppp cells in wild type (WT) (A), *tlk-1* mutants (B), *tlk-1 sin-3* double mutants (C), and *tlk-1 hira-1* double mutants (D) at the L3 stage. Arrows indicate the nucleus of V5.ppp. All images were captured by confocal microscopy at the same setting. Scale bars indicate 10 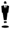 m. (E) Box-and-whisker plot showing the intensity of H3.3 (*his-72::gfp*) in the nucleus. *n* = 20. Whiskers indicate the 10th and 90th percentiles. Boxplots represent the medians and the 25th–75th percentile.

### Loss of SIN-3 suppresses extra-DTC phenotype of *tlk-1* mutants

We performed RNAi screening for suppressors of *tlk-1* mutants using the chromatin subset of the Ahringer’s feeding RNAi library (Kamath et al., 2003). We found that *sin-3* RNAi suppressed the extra-DTC phenotype of *tlk-1* mutants (data not shown). A *sin-3* deletion mutant also suppressed the *tlk-1* phenotype (Fig. 6A, Fig. S5A, B). Most *sin-3 tlk-1* double mutants had two DTCs. The frequency of animals with one or no DTCs was similar between the double mutants and the s*in-3* single mutants (Fig. S5A). Of note, *sin-3* RNAi also suppressed the extra-DTC phenotype of *chaf-1* mutants (Fig. 6B, Fig. S5C). Thus, *sin-3* was antagonistic to *tlk-1* rather than having a role in DTC differentiation.

**Fig. 6.**
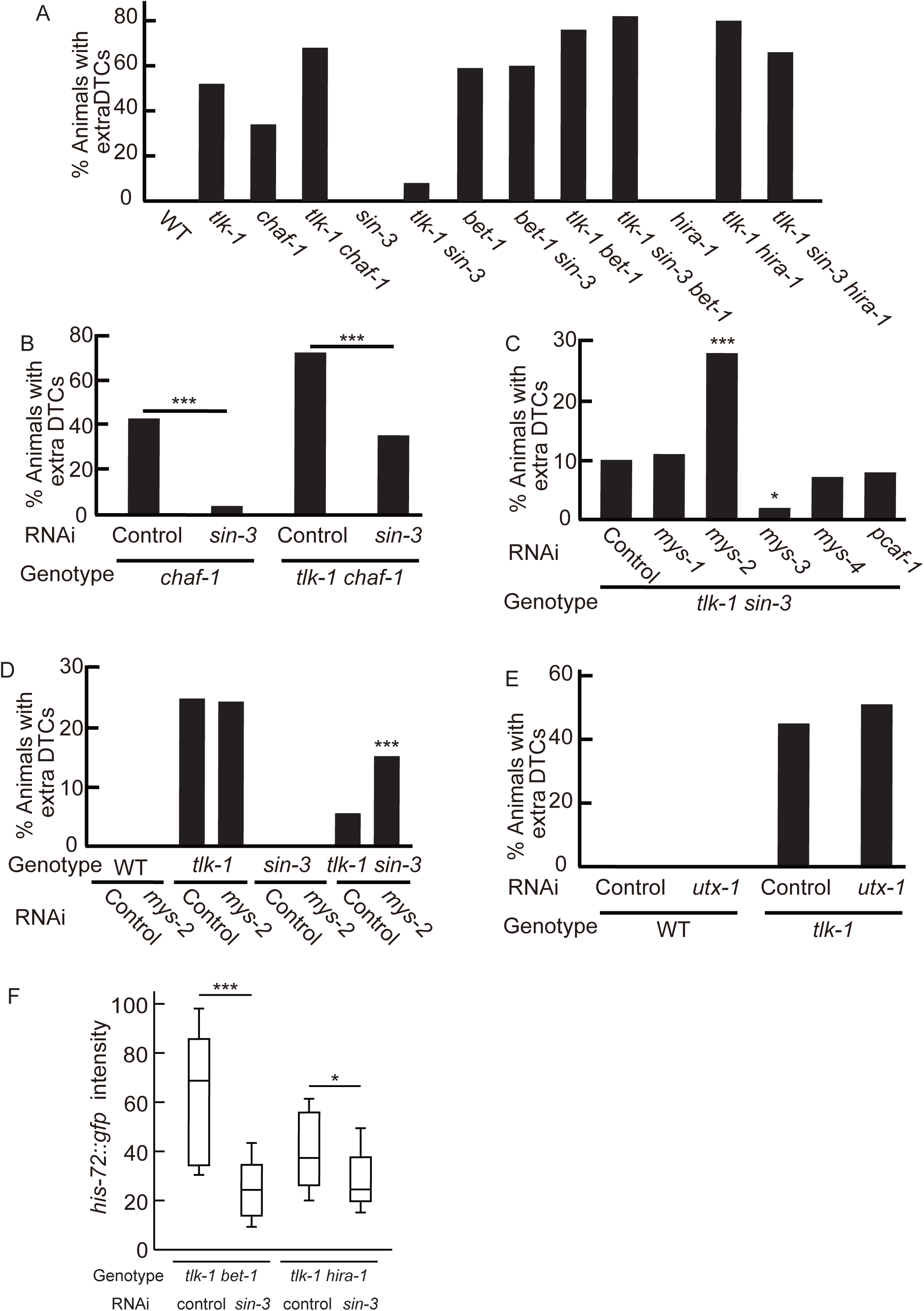
*sin-3* as a suppressor of *tlk-1*. (A–E) Bar graphs show the percent of adult animals with extra DTCs. *n* = 100. *p* values for (A) are indicated in Fig. S5B. For (B–E), *0.05 < *p* < 0.01, \*\*\**p* < 0.005, as compared with the control. (F) Box-and-whisker plot showing the intensity of H3.3 (*his-72::gfp*) in the nucleus. *n* = 20. Whiskers indicate the 10th and 90th percentiles. Boxplots represent the medians and the 25th–75th percentile. *0.05 < *p* < 0.01, \*\*\**p* < 0.005.

We also examined the expression of *his-72::gfp* in the *tlk-1 sin-3* background. Interestingly, the level of *his-72::gfp* expression also decreased in *tlk-1 sin-3* relative to its expression in *tlk-1* (Fig. 5C, E). There was a positive correlation between the *his-72::gfp* expression level and the extra-DTC phenotype. For example, half of *tlk-1* mutants and 8% of *tlk-1 sin-3* mutants showed the extra-DTC phenotype. The median *his-72::gfp* expression level in *tlk-1* mutants was almost the same as the 80^th^ percentile for *his-72::gfp* expression in *tlk-1 sin-3* mutants.

### AcH4-binding protein BET-1 is necessary for suppression by sin-3 disruption

The SIN3 complex contains histone deacetylase, HDAC (Hassig et al., 1997; Laherty et al., 1997). However, HDAC (*had-1, 2, 3, 4,* and *6*) RNAi did not suppress the extra-DTC phenotype in *tlk-1* mutants (data not shown). If histone hyper-acetylation in *tlk-1 sin-3* double mutants is responsible for suppression of the extra-DTC phenotype, disruption of histone acetyltransferase may induce extra DTCs. We found that RNAi of *mys-2*, which encodes MYST family histone acetyltransferase, induced the extra-DTC phenotype in the *tlk-1 sin-3* background (Fig. 6C, Fig. S5D). *mys-2* RNAi did not enhance the extra-DTC phenotype in the *tlk-1* background and did not induce extra DTCs in the wild-type or *sin-3* background (Fig. 6D, Fig. S5E). Therefore, the effect of *mys-2* RNAi was specific to the *tlk-1 sin-3* background. These results suggested that the extra-DTC phenotype in *tlk-1* mutants can be compensated for by hyper-acetylation.

MYS-2 controls sub-nuclear localization of BET-1 (Shibata et al., 2014). Because we originally isolated *tlk-1* mutants as a phenocopy of *bet-1* mutants, we examined the relationship between *tlk-1* and *bet-1*. First, we examined the phenotype of *tlk-1 bet-1* double mutants. We found that *tlk-1 (tk158)* enhanced the *bet-1* null allele, *os46* (Fig. 6A, Fig. S5A, B). The maximum number of DTCs was seven in *tlk-1 bet-1* double mutants, in contrast to four and five in *bet-1* and *tlk-1* mutants, respectively (Fig. S5A). Therefore, *tlk-1* acts in parallel with *bet-1* to maintain the DTC cell fate. We also found that the *tlk-1* suppressor *sin-3* could not suppress *bet-1* (Fig. 6A, Fig. S5A, B). The *bet-1* suppressor *utx-1* also could not suppress *tlk-1* (Fig. 6E, Fig. S5F). These results are consistent with parallel regulation by *bet-1* and *tlk-1*.

If SIN-3 regulates BET-1 through acetylation, *sin-3* suppression should be dependent on *bet-1*. However, if SIN-3 and BET-1 control parallel pathways, *sin-3* should suppress the extra-DTC phenotype of *tlk-1* mutants even without *bet-1*. The results revealed that *sin-3* did not suppress the extra-DTC phenotype of *tlk-1 bet-1* double mutants, indicating that the *sin-3* suppression was dependent on *bet-1* (Fig. 6A, Fig. S5A, B). Although the phenotypic penetrance of *tlk-1 chaf-1* was similar to that of *tlk-1 bet-1*, *sin-3* RNAi suppressed the extra-DTC phenotype only in the *tlk-1 chaf-1* background (Fig. 6B, Fig. S5C). These results indicated that *bet-1* is downstream of *sin-3*.

Next, we examined whether SIN-3 regulates H3.3 through BET-1. *sin-3* RNAi suppressed the level of *his-72::gfp* expression in *tlk-1 bet-1* background, indicating that BET-1 is not required for the suppression of H3.3 by *sin-3* depletion in *tlk-1* mutants. Thus, BET-1 is downstream of H3.3 accumulation.

### HIRA-1 depletion enhances *tlk-1* phenotypes

HIRA and Daxx are H3.3 chaperones in mammals (Lewis et al., 2010). *C. elegans* has a homolog of HIRA, HIRA-1, but it does not have a Daxx homolog. If HIRA-1 is a major H3.3 chaperone in *C. elegans*, *hira-1* mutations may suppress the extra-DTC phenotype of *tlk-1* mutants. However, unexpectedly, *hira-1* mutations enhanced the extra-DTC phenotype of *tlk-1* mutants (Fig. 6A, Fig. S5A, B). We also examined *his-72::gfp* expression and found that the *hira-1* mutation up-regulated the *his-72::gfp* level in the wild-type and *tlk-1* backgrounds (Fig. 5D, E). Because the enhancement of the extra-DTC phenotype by the *hira-1* mutation was similar to that of the *bet-1* or *chaf-1* mutations, we examined whether *sin-3* could suppress the extra-DTC phenotype in *tlk-1 hira-1* mutants. The results revealed that the *sin-3* mutation showed weak suppression in *tlk-1 hira-1* mutants (Fig. 6A, Fig. S5A, B). These results suggested that the functions of HIRA proteins may have diverged between *C. elegans* and mammals.

We also examined whether *hira-1* is required for down-regulation of *his-72::gfp* by *sin-3* depletion. *sin-3* RNAi suppressed the level of *his-72::gfp* expression in *tlk-1 hira-1* background. However, compared to *tlk-1* or *tlk-1 bet-1* background, the suppression was weak. This result indicated that the down-regulation of *his-72::gfp* by *sin-3* depletion is partially dependent on *hira-1*.

## Discussion

### TLK-1 and CHAF-1 maintain cell fates

In this paper, we showed that TLK-1 and CHAF-1 maintain the cell fates of multiple cell types by inhibiting their fate transformation into gonadal DTCs, PDE and PVD interneurons, and AVM and PVM touch receptor neurons. TLK-1 represses, directly or indirectly, *ceh-22* and *mec-3*, which encode the NK-2 family homeodomain and LIM homeodomain DNA-binding TFs, respectively. We speculate that TLK-1 regulates transcription of the selector genes that encode DNA-binding TFs (Garcia-Bellido, 1975; Hobert, 2008). Although it is not clear whether the *mec-3* locus is a direct target of TLK-1, ectopic production of *mec-3*-expressing cells in *tlk-1* mutants and the loss of *mec-4* expression in *tlk-1 mec-3* double mutants suggest this possibility. The reduction of ectopic DTCs by *ceh-22* RNAi in *tlk-1* mutants also suggests that *ceh-22* is a functional target.

### TLK-1 and CHAF-1 regulate histone H3.3 in cell-fate maintenance

Disruption of TLK-1 or CHAF-1 has the same effect in terms of extra DTCs, extra PDEs, extra AVMs, extra PVMs, and higher histone H3.3 expression. Together with biochemical evidence in other organisms, TLK-1 also appears to function with the CAF1 complex in *C. elegans*. The CAF1 complex deposits the histone H3-H4 complex as a DNA replication–coupled histone chaperone (Klimovskaia et al., 2014). The levels of H3.3 were elevated in *tlk-1* and *chaf-1* mutants. Interestingly, the expression level of H3.3 correlated with the penetrance of the extra-DTC phenotype, suggesting that H3.3 plays an important role in cell-fate maintenance. We speculate that the failure of H3 deposition because of the loss of *chaf-1* activity induces recruitment of H3.3, which could substitute for H3 in the essential function of the nucleosomes. Because H3.3 localizes on actively transcribed loci (Wirbelauer et al., 2005), H3.3 appears to promote transcriptional activation. It is not clear whether loss of H3 or ectopic deposition of H3.3 is responsible for the defect in cell-fate maintenance in *tlk-1* and *chaf-1* mutants.

### TLK-1 and BET-1 act in parallel pathways

We found that loss of SIN-3 strongly suppressed the extra-DTC phenotype of *tlk-1* and *chaf-1* mutants. The SIN3 complex contains HDAC (Hassig et al., 1997; Laherty et al., 1997), suggesting that the acetylation of histones is elevated in *sin-3* mutants. As expected, RNAi knockdown of one of the histone acetyl transferases, *mys-2*, reversed the suppression effect of *sin-3* mutants. Furthermore, the suppressor activity of *sin-3* mutants depended on an acetylated histone H4 binding protein, BET-1, required for fate maintenance of cell types whose fates are also maintained by TLK-1 and CHAF-1 (Shibata et al., 2014). Because the cell fate-maintenance defect was enhanced in *tlk-1* and *bet-1* double mutants, it is likely that TLK-1 and BET-1 act in distinct pathways to regulate cell-fate maintenance.

We previously showed that the BET-1 pathway represses transcription of target genes by recruiting H2A.z to maintain cell fates (Shibata et al., 2014). Loss of H2A.z is the major cause of defects in cell-fate maintenance in *bet-1* mutants (Fig. S6B). In contrast, repression of the extra-DTC phenotype in *tlk-1 sin-3* double mutants appears to be caused by an H3-dependent mechanism, because there is a positive correlation between the level of H3.3 and the strength of the extra-DTC phenotype in *tlk-1 sin-3* double mutants, *tlk-1* mutants, *chaf-1* mutants, and *tlk-1 hira-1* double mutants. Therefore, it is likely that the TLK-1 and CHAF-1 pathway negatively regulates H3.3 and that the SIN-3, MYS-2, and BET-1 pathway positively regulates H2A.z deposition.

### Model for cell-fate maintenance by histone variants

Based on our findings, we propose the following model. Localization of histone H3 on selector gene loci is critical for transcriptional repression (Fig. 7A). H3 deposition by TLK-1 and CAF1 inhibits deposition of H3.3. In *tlk-1* mutants, reduction of CAF1 activity causes ectopic H3.3 deposition when the SIN-3-HDAC complex deacetylates histone H4 (Fig. 7B). Stochastic expression of selector genes by ectopic H3.3 deposition causes cell-fate transformation. In *tlk-1 sin-3* double mutants, loss *sin-3* activates two independent downstream machineries. One is a HIRA-1-dependent mechanism which causes down-regulation of H3.3 (Fig. 7C). Although HIRA is known as H3.3 chaperone in mammal, *C. elegans* HIRA-1 may promote H3 deposition. The other mechanism is activated by hyper-acetylation of H4 that is caused by malfunction of the SIN-3/HDAC complex. Hyper-acetylation of H4 is recognized by BET-1 that promotes H2A.z incorporation. BET-1-dependent mechanism does not affect the level of H3.3. Thus, loss of *sin-3* causes down-regulation of H3.3 and deposition of H2A.z in a manner dependent on HIRA-1 and BET-1, respectively.

**Fig. 7.**
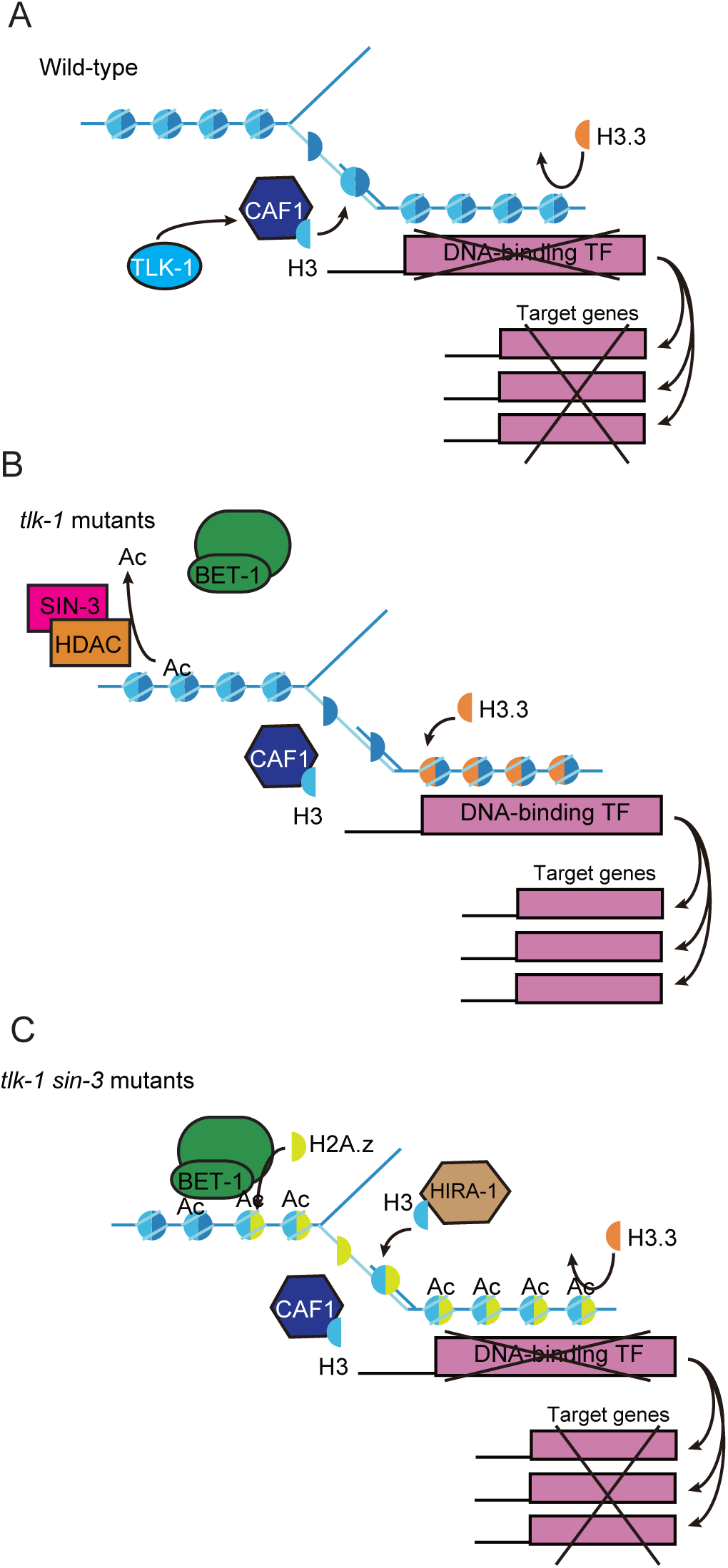
Model for cell-fate maintenance by histone variants. (A) In wild-type animals, TLK-1 and CAF1 promote formation of H3-containing nucleosomes that prevent aberrant deposition of H3.3. (B) In *tlk-1* mutants, H3.3 is incorporated into the nucleosome-free region that is formed by dysfunction of the CAF1 complex. H3.3 causes stochastic expression of DNA-binding TFs. (C) In *tlk-1 sin-3* double mutants, hyper-acetylation of histones promotes recruitment of the BET-1 complex, which then forms H2A.z-containing nucleosomes. Loss of *sin-3* also decreases H3.3 in a manner dependent on HIRA-1. HIRA-1 may promote H3 deposition.

Although it is thought that H2A.z localization is correlated with transcriptional activation, we previously showed that H2A.z represses transcription in cell-fate maintenance (Shibata et al., 2014). These results, together with the current research, lead us to speculate that H3.3 localization is correlated with transcriptional activation, whereas H2A.z localization is correlated with transcriptional repression in cell-fate maintenance. In agreement, H2A.z-mediated chromatin compaction is repressed by H3.3 *in vitro* (Chen et al., 2013). These histone variants may co-regulate transcription through chromatin compaction in cell-fate maintenance. Chromatin compaction may affect the binding of TFs to cis elements, association of transcriptional machinery, or transcriptional elongation by RNA pol II.

Although HIRA is known as a chaperone of H3.3 in mammals (Tagami et al., 2004), a *hira-1* deletion up-regulated the H3.3 level in *C. elegans*. One explanation is that *C. elegans* HIRA-1 function as a chaperone of H3. Alternatively, this may be explained by the activation of another H3.3 chaperone, for example, the counterpart of Daxx (Lewis et al., 2010). However, there is no Daxx homolog Daxx in *C. elegans*. *C. elegans* may have other H3.3 chaperones. It is reported that knockdown of the H2A.z chaperone EP400 hampers H3.3 deposition in a human cell line (Pradhan et al., 2016). However, a similar mechanism is inconsistent with down-regulation of HIS-72::GFP by activation of BET-1, which promotes H2A.z deposition. Thus, it is likely that the function of *C. elegans* HIRA-1 may have diverged from HIRA proteins of other organisms during evolution.

Recent studies revealed that artificial induction of trans-differentiation, including generation of iPS cells, is improved by repression of CAF1 (Cheloufi et al., 2015), which is known as a downstream factor of TLK (Klimovskaia et al., 2014). The current study is the first example showing the role of CAF1 and TLK-1 in cell-fate maintenance during normal development. The roles of CAF1, and probably TLK, in the maintenance of cell fate appear to be conserved in multicellular organisms. Our research also showed that there is a strong correlation between the H3.3 level and defects in cell-fate maintenance, suggesting that the regulation of H3 variants is important for cell-fate maintenance and trans-differentiation. We also unveiled the regulation of H3 variants through BET-1, which promotes H2A.z deposition. H3.3 and H2A.z are major histone variants that are conserved in yeast, *C. elegans*, and mammals. Crosstalk between these histone variants may be a fundamental mechanism in the repression of trans-differentiation. The roles of H3.3, BET family proteins, and H2A.z remain unknown in artificial trans-differentiation. Thus, the functional conservation of TLK, CAF1, and histone variants between cell-fate maintenance in *C. elegans* and artificial trans-differentiation in mammals is a compelling issue for future studies.

## Materials and Methods

### Strains and culture

N2 Bristol was used as the wild-type *C. elegans* strain (Brenner, 1974). Animals were cultured at 20°C. The *bet-1* (Shibata et al., 2010)*, chaf-1, rba-1* (Nakano et al., 2011), and *tlk-1* mutants are sterile and were maintained as heterozygotes over the hT2[qIs48] balancer. The phenotypes of homozygotes generated from the heterozygous hermaphrodites were analyzed. The following green fluorescent protein (GFP) and red fluorescent protein (RFP) markers were used: *zdIs5[mec-4::gfp]* (Clark and Chiu, 2003), *qIs56[lag-2::gfp]* (Kostic et al., 2003), *mnIs17[osm-6::gfp]* (Collet et al., 1998), *vsIs33[dop-3::rfp]* (Chase et al., 2004), *uIs22[mec-3::gfp]* (Toker et al., 2003), *vtIs1[dat-1::gfp]* (Nass et al., 2002), and *stIs10026[his-72::gfp]* (Boeck et al., 2011). Synchronization of animals was performed as described (Shibata et al., 2010).

### RNAi

The *sin-3*, *mys-1*, *mys-2*, *mys-3*, *mys-4*, *pcaf-1*, *hda-1*, *hda-2*, *hda-3*, *hda-4*, *hda-6*, and *ceh-22* RNAi constructs were described previously (Kamath et al., 2003; Shibata et al., 2010). Feeding RNAi experiments were performed as described (Kamath et al., 2001). RNAi screening was performed using the *C. elegans* RNAi chromatin library (Source BioScience, Nottingham, UK).

### Cloning of tlk-1

Single-nucleotide polymorphism (SNP) mapping indicated that both *tk158* and *tk170* are positioned on LG III. A complementation test revealed that *tk158* and *tk170* are allelic (data not shown). Because *tk158* showed a more severe phenotype (Fig. 1E, Fig. S1A), we used *tk158* for further analysis. *tk158* was positioned at the center cluster of LGIII between 0.92 and 1.13 by SNP mapping (Fig. 1F). Because *tk158* and *tk170* are sterile, heterozygous animals that are balanced by hT2 were used for genome sequencing. Within the candidate region, *tlk-1* is the sole gene that has a non-synonymous mutation in both *tk158* and *tk170* mutants. For the rescue experiment, a PCR fragment that contained *tlk-1* and 3.5 kb of upstream sequence was amplified from the fosmid WRM0631bB10 using primers 5′-CTCTCTTTGCCACTTTATCGTTTGT-3′ and 5′-AAGTTTGCGCATGTAGTAAGTTTCA-3′. The transgenic marker was *myo-3::mCherry*.

### Microscopy and statistical analysis

Expression of *lag-2::gfp*, *dat-1::gfp*, *osm-6::gfp*, *dop-3::rfp*, and *mec-3::gfp* was detected by epifluorescence microscopy (AxiosImagerM2 and Axioplan2; Zeiss, Jena, Germany). Expression of *his-72::gfp* was detected by confocal microscopy (LSM510 and Pascal; Zeiss) in the L3 animals. We used V5.ppp to quantify *his-72::gfp* because V5.ppp is easy to identify in wild-type animals and mutants, has a large nucleus, and is positioned near the body surface. The average intensity in the V5.ppp nucleus was measured using ImageJ (National Institutes of Health, Bethesda, MD). Background fluorescence was measured from the adjacent region of the nucleus of V5.ppp and subtracted from the average intensity.

## Acknowledgements

We thank the Caenorhabditis Genetics Center, which is funded by the National Institutes of Health National Center for Research Resources, and the National Bioresource Project for strains. This work was supported by grants from the Japanese Ministry of Education, Culture, Sports, Science and Technology (Y.S. and K.N.).

